# mock-fastq-generator: A synthetic FASTQ generator

**DOI:** 10.64898/2026.07.28.740902

**Authors:** Alberto Florez Prada, Darren J. Hart

## Abstract

Validating bioinformatics pipelines and benchmarking sequence processing algorithms requires reliable test datasets. Existing read simulation tools rely on reference genomes and empirical error profiles, lacking fine-grained control over specific targeted DNA constructs and controlled error injection. `mock-fastq-generator` is an open-source software suite available both as an installable PyPI Python package and a standalone, client-side web application. It constructs synthetic FASTQ files by combining template constructs with customizable adapter sequences, selectable quality decay functions (Gaussian, Exponential, Sigmoidal), NovaSeq 3-state quality binning, and context-dependent error penalties. The software allows developers to benchmark sequence trimmers, test alignment sensitivity, and execute automated quality control pipelines in test suites without using proprietary or empirical biological data.

## 1. Motivation and significance

Next-generation sequencing (NGS) workflows depend on computational software for quality filtering, adapter trimming, read alignment, and variant identification. Developing and testing these analytical pipelines requires reliable test datasets. Utilizing empirical sequencing data during software development introduces confounding biological variability, limited capacity for testing edge cases, and potential intellectual property or privacy concerns.

Thus, synthetic read generation offers a controlled environment to evaluate algorithmic behavior.

Traditional read simulation software packages, such as ArtificialFastqGenerator [1], ART [2], and DWGSIM [5], construct simulated reads by sampling reference genomes according to empirical error distributions. However, deploying complex software overcomplicates a simple task with unnecessary deployment-related technical demands. Other specialized software solutions target third-generation long-read platforms [3] or single-cell transcriptomics [4]. Empirical error profilers, including GemSIM [7], capture technology-dependent error profiles to replicate platform behaviors [8]. While effective for genome-wide evaluation, these frameworks require full reference genomes and lack mechanism-level control over localized quality degradation, making them unsuitable for targeted construct simulation or isolated unit testing. Conversely, basic inline scripts that assign uniform quality scores, such as constant Q40 values, fail to represent realistic physical error distributions.

`mock-fastq-generator` addresses the requirement for lightweight, targeted synthetic data generation. The software combines construct-level sequence assembly with customizable mathematical quality decay functions, context-dependent error penalties, and discrete quality binning. The tool enables developers to inject controlled quality drops at specified sequence coordinates, benchmark pipeline components, and validate bioinformatics software deterministically.

## 2. Software description

`mock-fastq-generator` is implemented in Python and native JavaScript, structured for modular execution, script automation, and browser-based interactive exploration.

### 2.1. Software Architecture and Quality Assurance

The software architecture consists of two implementations. The primary Python package follows standard distribution practices and is installable directly via PyPI (pip install mock−fastq−generator). Automated unit testing via `pytest` achieves 97% statement coverage across module functions. Continuous Integration (CI) and automated regression testing upon codebase modifications are managed using GitHub Actions workflows.

Complementing the Python distribution, a client-side web application was built with standard HTML5, CSS3, and JavaScript (https://florez-alberto.github.io/mock-fastq-generator/). The web interface executes all sequence assembly and quality score calculations locally within the browser, integrating real-time visualization of positional Phred score decay curves and client-side file generation without external server dependencies.

The core pipeline architecture contains two principal components: the Sequence Assembly Module and the Quality Modeling Module (Figure 1).

**Figure 1:**
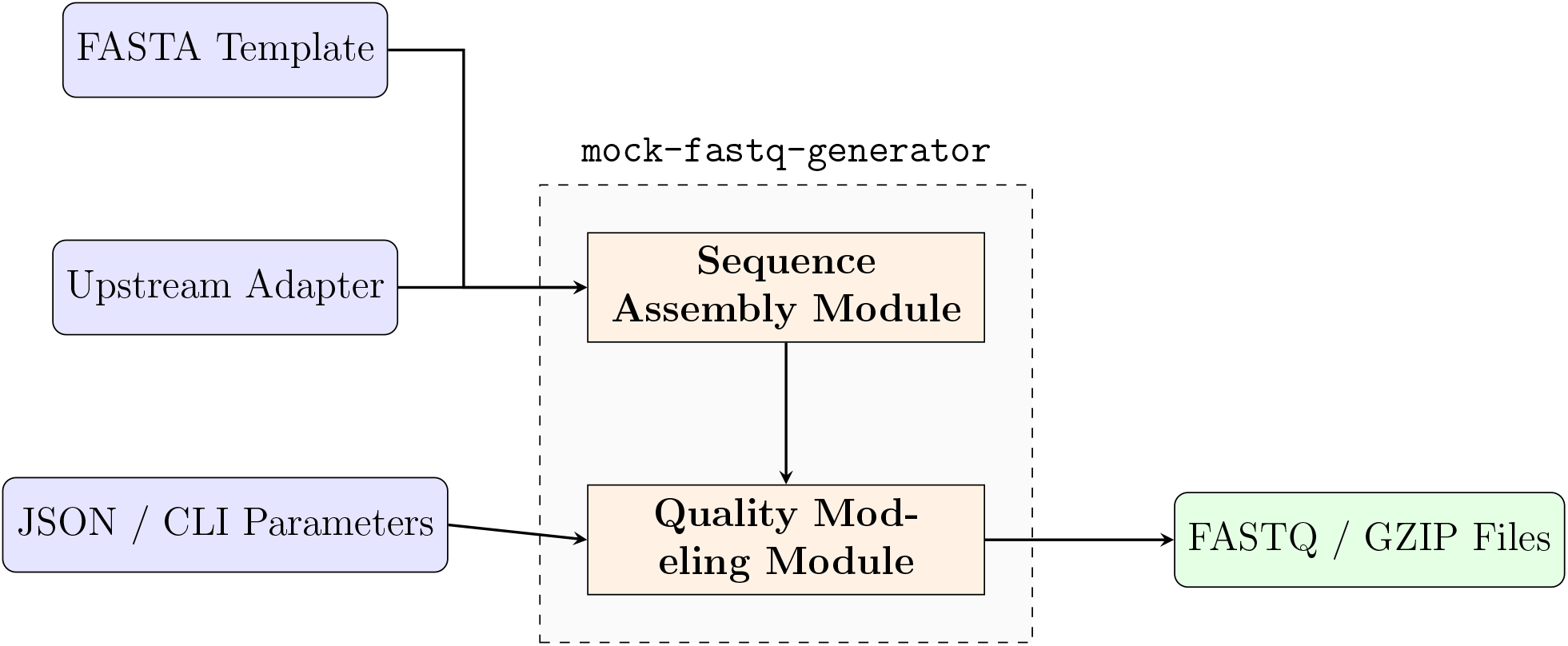
System architecture and operational data flow of `mock-fastq-generator`. Target template sequences and configuration options are processed by the sequence assembly and quality modeling modules to produce standardized FASTQ streams or compressed files.

The Sequence Assembly Module parses user-supplied FASTA template records, prepends a defined upstream sequence, and inserts randomized nucleotide margins to construct full-length synthetic reads. Substitution noise is introduced into the template region according to a specified mutation probability.

The Quality Modeling Module evaluates the assembled sequence to generate positional Phred quality scores. Generated continuous scores are clamped to defined boundaries and converted to ASCII characters adhering to the standard Sanger Phred offset (+33) [11].

### 2.2. Core Functionalities and Performance Metrics

The software supports comprehensive configuration options for data generation:

- **Single-end and Paired-end Modes:** Paired-end mode (`--paired-end`) constructs matching forward (R1) and reverse complement (R2) files with designated read-pair suffixes.
- **Output Stream Flexibility:** Supports plain text FASTQ, inline gzip compression (`--gzip`), and direct standard output streaming (`--stdout`) to pipe data directly into aligners without disk I/O bottlenecks.
- **Configuration Repositories:** CLI parameters or structured JSON configuration files ensure reproducible dataset generation across automated execution environments.

Throughput performance benchmarks conducted on a standard workstation demonstrate generation speeds of 0.11 s for 1,000 reads (∼9,100 reads/s), 1.01 s for 10,000 reads (∼9,900 reads/s), and 10.08 s for 100,000 reads (∼9,900 reads/s).

### 2.3. Mathematical Quality Decay and Error Models

Let *i* ∈ {0, 1, …, *N* −1} represent the zero-indexed nucleotide position within a read of length *N* . The unconstrained quality value *Q*_raw_(*i*) is computed according to the selected mathematical model:

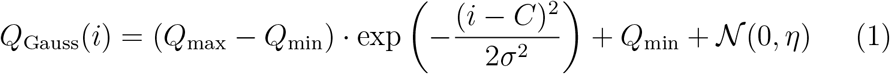

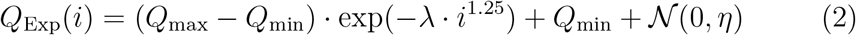

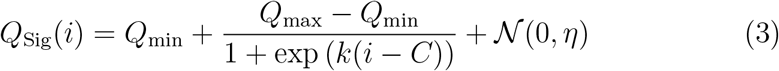

where *Q*_min_ and *Q*_max_ define quality boundaries, *C* denotes the inflection or peak center index, *σ* specifies standard deviation, *λ* represents the exponential decay rate, *k* dictates sigmoidal steepness, and *N*(0, *η*) adds Gaussian noise. The fractional exponent 1.25 in the exponential model introduces a sub-exponential initial plateau representing early high-quality cycles.

A context-dependent homopolymer degradation rule applies a penalty Δ_*H*_ = 10 for any position *i* within runs of > 4 identical nucleotides:

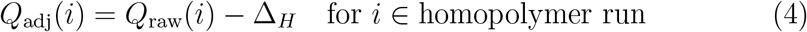

Adjusted continuous scores are clamped to [*Q*_min_, *Q*_max_]. When 3-state NovaSeq quality binning is enabled (emulating modern NovaSeq X RTA4 chemistry), scores are mapped to discrete bins *b* ∈ ℬ = {12, 24, 40} (or custom user-defined bin sets such as {12, 23, 37} for NovaSeq 6000 RTA3):

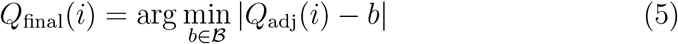

## 3. Illustrative examples

The CLI tool is invoked via `mock-fastq-generator`. To generate a dataset with a 3^*′*^ sigmoidal quality drop centered at position *C* = 100 (*k* = 1.0) for sliding-window trimming benchmarks:

~~~
mock-fastq-generator \
 --output_file sigmoidal_benchmark.fastq \
 --template_sequence target.fasta \
 --total_length 150 \
 --number_of_sequences 100 \
 --decay_model sigmoidal \
 --decay_rate 1.0 \
 --center 100 \
 --min_val 5 \
 --max_val 40 \
 --score_type phred
~~~

To generate a paired-end dataset featuring 3-state NovaSeq X quality score binning and homopolymer penalties:

~~~
mock-fastq-generator \
 --output_file paired_sample \
 --template_sequence target.fasta \
 --paired-end \
 --number_of_sequences 100 \
 --binned_quality \
 --homopolymer_penalty
~~~

For streaming directly into processing pipelines:

~~~
mock-fastq-generator \
 --template_sequence target.fasta \
 --number_of_sequences 500 \
 --stdout | bwa mem reference.fasta - > mapped_reads.sam
~~~

Figure 2 displays the generated decay profiles.

**Figure 2:**
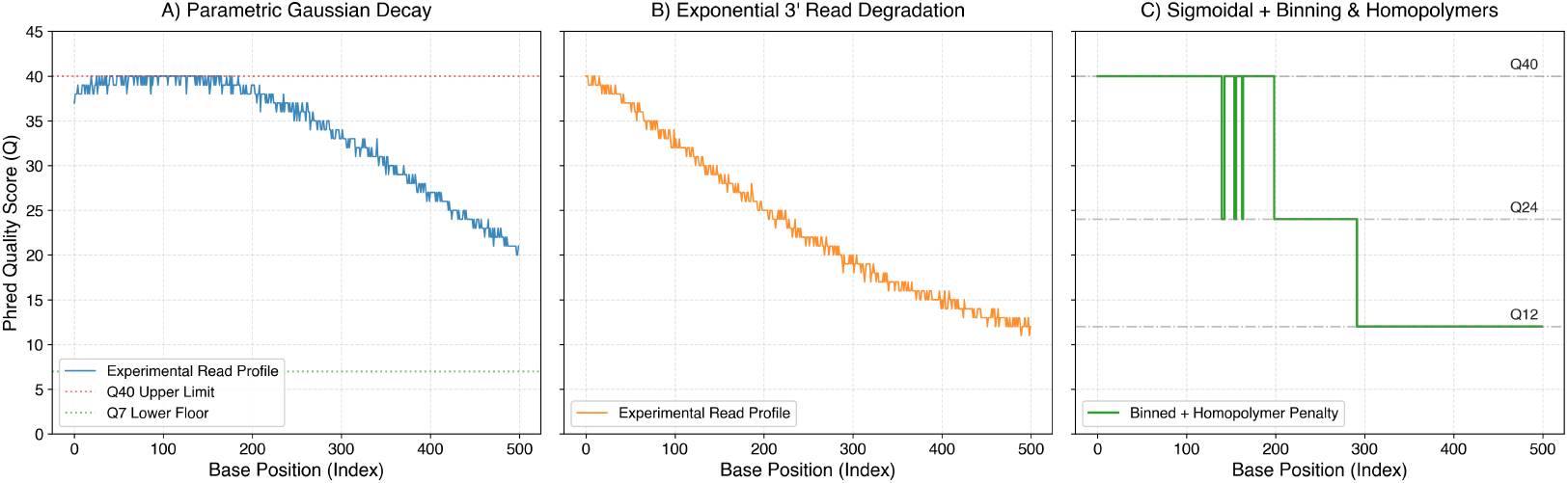
Comparison of synthetic Phred quality score degradation profiles generated across a 500-base sequence construct. (A) Parametric Gaussian decay (*C* = 100, *σ* = 300). (B) Exponential decay modeling 3^*′*^ quality dropoff. (C) Sigmoidal drop combined with 3-state Illumina NovaSeq X binning (*Q*12, *Q*24, *Q*40) and homopolymer penalties.

## 4. Impact

`mock-fastq-generator` provides a controlled environment for evaluating read preprocessing software [13, 12, 14], amplicon denoising algorithms [9], and sequence clusterers [10].

To demonstrate utility for quantitative pipeline validation, a benchmark experiment was conducted using `fastp` [12] (Table 2). Synthetic read datasets (*N* = 100, 150 bp) were generated under three controlled conditions. When evaluated against the standard sliding-window quality threshold of `fastp` (*Q* ≥ 15), low-quality reads were 100% rejected, high-quality controls were 100% retained, and sigmoidal drop reads were truncated exactly at position 100 ± 1 bp.

**Table 1:**
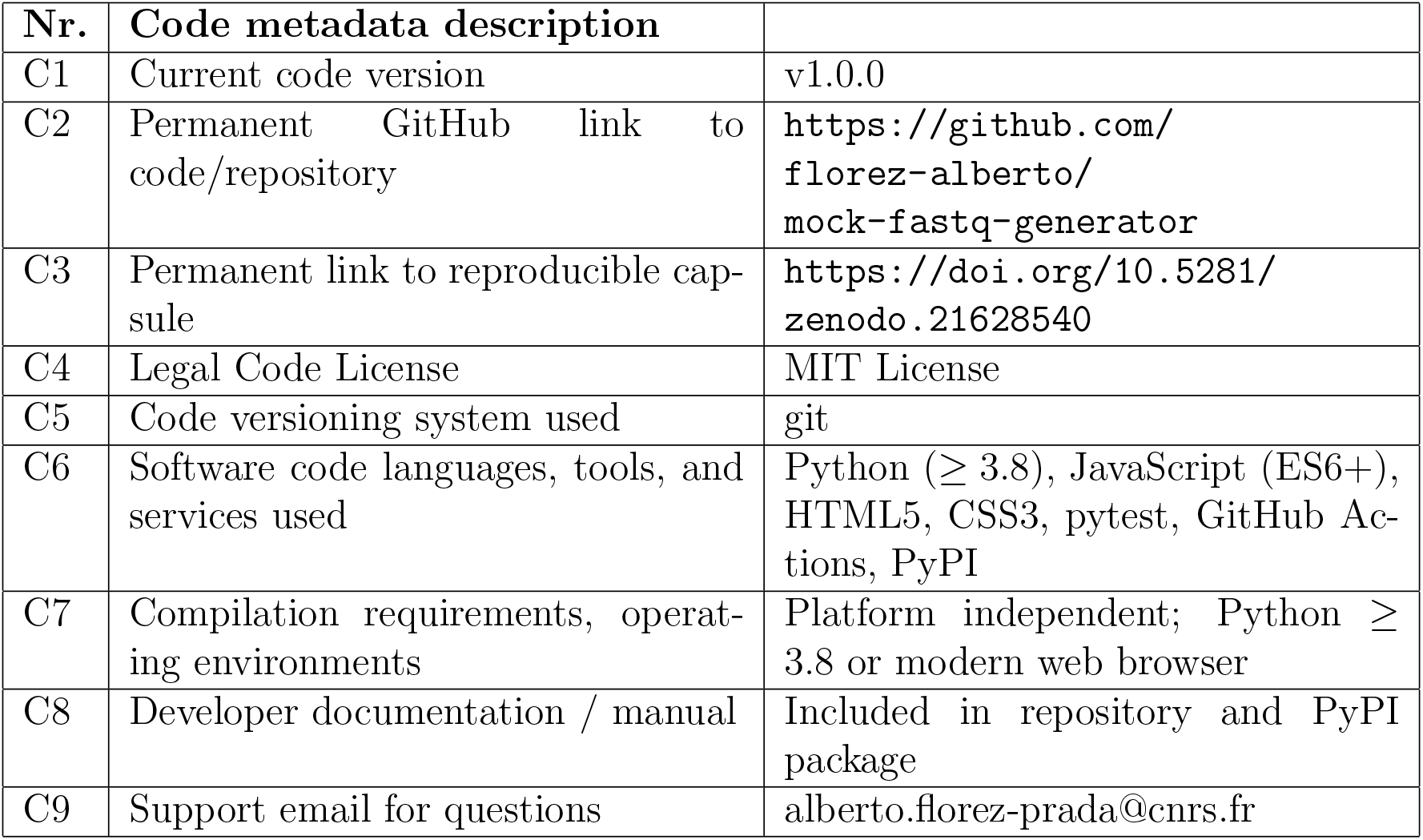
Code metadata.

**Table 2:** Quantitative benchmark evaluating fastp preprocessing behavior using synthetic FASTQ datasets generated under controlled quality conditions.

| Synthetic Condition | Quality Range ($Q$) | fastp Output Result | Filter Efficiency |
| --- | --- | --- | --- |
| Low-Quality | $Q \in [0, 7]$ ($Q < 15$) | 100/100 reads rejected | 100% low-quality filter |
| High-Quality Control | $Q \in [27, 40]$ ($Q \geq 15$) | 100/100 reads retained | 100% pass-through |
| 3' Sigmoidal Drop ($C = 100$) | $Q_{40} \rightarrow Q_5$ drop at pos 100 | Trimmed at pos $100 \pm 1$ | Exact boundary truncation |

## 5. Conclusions

`mock-fastq-generator` provides a lightweight, deterministic framework for generating synthetic FASTQ sequencing data based on construct-based assembly and mathematical decay models. Available as both a PyPI Python package and a zero-dependency web interface, it establishes a standardized platform for automated unit testing and quality control pipeline validation.

## 6. Declarations

### CRediT authorship contribution statement

**Alberto Florez Prada:** Conceptualization, Methodology, Software, Validation, Formal analysis, Investigation, Writing – original draft, Writing – review & editing. **Darren J. Hart:** Conceptualization, Supervision, Funding acquisition, Writing – review & editing.

## Acknowledgements

The authors acknowledge support from the Centre National de la Recherche Scientifique (CNRS). AFP and DH received funding from the ANR FluPept project ANR-21-CE18-0024 and the France 2030 PUI program of Université Grenoble Alpes (UGA). IBS acknowledges integration into the Interdisciplinary Research Institute of Grenoble (IRIG, CEA). AFP received funding from the École doctorale Chimie et sciences du vivant (EDCSV) – UGA.

## Declaration of generative AI in manuscript preparation

During the preparation of this work, the authors used AI tools for text clarity, LaTeX formatting, and code style adherence. The authors reviewed and edited the content as needed and take full responsibility for the published article.

## References

[1] M. Frampton, R. Houlston, Generation of artificial FASTQ files to evaluate the performance of next-generation sequencing pipelines, PLoS ONE 7(10) (2012) e49110.

[2] W. Huang, L. Li, J.R. Myers, G.T. Marth, ART: a next-generation sequencing read simulator, Bioinformatics 28(4) (2012) 593–594.

[3] Y. Ono, K. Asai, M. Hamada, PBSIM3: a simulator for all types of PacBio and ONT long reads, Bioinformatics 38(5) (2022) 1415–1417.

[4] G. Baruzzo, I. Vurchio, C. Romualdi, B. Di Camillo, SPARSim single cell: a count data simulator for scRNA-seq data, Bioinformatics 36(5) (2020) 1468–1475.

[5] N. Homer, DWGSIM: A whole genome simulator for next-generation sequencing, https://github.com/nh13/DWGSIM (2010).

[6] A. Bennett, C.J. Miller, L. Huet, P. Bayer, nf-core/readsimulator: A pipeline to simulate sequencing reads, Zenodo (2024) doi:10.5281/zenodo.10622410.

[7] K.E. McElroy, F. Luciani, T. Thomas, GemSIM: general, error-model based simulator of next-generation sequencing data, BMC Genomics 13(1) (2012) 74.

[8] M. Schirmer, R. D’Amore, U.Z. Ijaz, N. Hall, C. Quince, Illumina error profiles: resolving fine-scale variation in metagenomic sequencing data, BMC Bioinformatics 17(1) (2016) 125.

[9] B.J. Callahan, P.J. McMurdie, M.J. Rosen, A.W. Han, A.J.A. Johnson, S.P. Holmes, DADA2: High-resolution sample inference from Illumina amplicon data, Nature Methods 13(7) (2016) 581–583.

[10] R. Müller, M. Nebel, On the use of sequence-quality information in OTU clustering, PeerJ 9 (2021) e11717.

[11] P.J. Cock, C.J. Fields, N. Goto, M.L. Heuer, P.M. Rice, The Sanger FASTQ file format for sequences with quality scores, and the Solexa/Illumina FASTQ variants, Nucleic Acids Research 38(6) (2010) 1767–1771.

[12] S. Chen, Y. Zhou, Y. Chen, J. Gu, fastp: an ultra-fast all-in-one FASTQ preprocessor, Bioinformatics 34(17) (2018) i884–i890.

[13] A.M. Bolger, M. Lohse, B. Usadel, Trimmomatic: a flexible trimmer for Illumina sequence data, Bioinformatics 30(15) (2014) 2114–2120.

[14] M. Martin, Cutadapt removes adapter sequences from high-throughput sequencing reads, EMBnet.journal 17(1) (2011) 10–12.

